# Stool methanogens in intestine mammal species

**DOI:** 10.1101/2020.08.24.264788

**Authors:** C.O. Guindo, B Davoust, M Drancourt, G Grine

## Abstract

Methanogens are being members of anaerobe microbiota of the digestive tract of both human and mammals. However, the sources, modes of acquisition and dynamics of digestive tract methanogens remain poorly investigated. In this study, we aimed to expand the spectrum of animals which could be sources of methanogens for human, by exploring methanogen carriage in animals in contact with the general population or with some restricted populations; comparing the repertoire of animal methanogens with the one of human methanogens in order to question methanogens as zoonotic microorganisms. We used RT-PCR, PCR-sequencing and multispacer sequence typing to investigate the presence of methanogens in 407 fecal specimens collected from nine different mammalian species. We detected by RT-PCR, the presence of methanogen DNA in all mammals here investigated and none of the negative controls. We obtained by sequencing, seven different species of methanogens, of which three (Methanobrevibacter smithii, Methanobrevibacter millerae and Methanomassiliicoccus luminyensis) are known to be part of the methanogens present in the human digestive tract. We obtained 24 M. smithii by PCR-sequencing including 12/24 (50%) in pigs, 6/24 (25%) in dogs, 4/24 (16.66%) in cats, and 1/24 (4.16%) in both sheep and horses. Genotyping these 24 M. smithii revealed five different genotypes, all know in humans. Our results are fairly representative of the methanogen community present in the digestive tract of certain animals domesticated by humans and other future studies must be done to try to cultivate methanogens here detected by molecular biology to better understand the dynamics of methanogens in animals and also the likely acquisition of methanogens in humans through direct contact with these animals or through consumption of the meat and/or milk of certain animals, in particular cows.

## INTRODUCTION

Methanogens are archaea characterized by their unique capability in producing methane from by-products of bacterial anaerobe fermentations; being members of anaerobe microbiota of the digestive tract microbiota of several mammals (1). Accordingly, methanogens gained interest in the clinical microbiology over the last years after methanogens have been detected by PCR-based methods and cultured from the gut microbiota (2,3); and their translocation in milk and urines has been further observed (4). Moreover, methanogens have been associated with dysbiosis such as in the case of vaginosis (5), urinary tract infections (6) and anaerobe abscesses of the oral cavity in the case of periodontitis and periimplantitis (7,8), in the case of refractory sinusitis (9), brain (10,11) and muscle (12). Recently, we observed blood-borne methanogens associated with endocarditis (13). In all these situations, anaerobe bacteria were associated in the methanogen disease process and this observation was probably reflecting methanogen specificities including the absolute oxygen intolerance and the necessity of a hydrogen source to produce methane (14,15).

Currently, 16 different methanogens have been cultured from digestive tract microbiota of mammals (16–19) and PCR-based methods of detecting species-specific sequences traced an additional 04 species (18) (Table 1).

**Table 1.** Methanogens found in digestive tract microbiota of mammals.

| Culture | Additional species detected by PCR- based methods |
| --- | --- |
| *Methanosarcina* sp., *Methanobacterium formicicum*, *Methanomicrobium mobile*, *Methanosarcina barkeri*, *Methanobacterium bryantii*, *Methanobrevibacter ruminantium*, *Methanobrevibacter millerae*, *Methanobrevibacter olleyae*, *Methanoculleus olentangyi*, *Methanobrevibacter woesei*, *Methanobrevibacter gottschalkii*, *Methanobrevibacter thaueri*, *Methanobrevibacter wolinii*, *Methanobrevibacter cuticularis*, *Methanobrevibacter curvatus*, *Methanobrevibacter filiformis* | *M. smithii*, *Methanimicrococcus* spp., *Methanosphaera* spp., *Methanobacterium* spp. |

The sources, modes of acquisition and dynamics of digestive tract methanogens remain poorly investigated. We previously reported that the one-day newborns exhibited culturable *Methanobrevibacter smithii* (*M. smithii*) in the gastric fluid (20), suggesting a perinatal source of acquisition. Accordingly, we reported that mother milk did contain culturable *M. smithii* and culturable *Methanobrevibacter oralis* (*M. oralis*) (4). Yet, it is unclear whether these one-day methanogens do persist along with the digestive tract of the newborns or whether this is just one of several waves of acquisition of methanogens along the first months of the life (20–24). Therefore, the search for methanogens sources other than the mother milk is of interest.

Certain mammals (cow, sheep, donkey, horse, cat, pig, rabbit, rat, rhinoceros, baboon, monkey, and hippopotamus); birds (goose, turkey, and chicken) and insects (termites) are acknowledged to harbor digestive tract methanogens and *M. smithii* in particular has already been detected from bovine and also from Wistar rats (18,19,25–40). In this study, we aimed to expand the spectrum of animals which could be sources of methanogens for human, by exploring methanogen carriage in animals in contact with the general population or with some restricted populations; comparing the repertoire of animal methanogens with the one of human methanogens in order to question methanogens as zoonotic microorganisms.

## MATERIALS AND METHODS

### Feces samples

Feces are legally regarded as garbage in FRANCE and their collection does not require any institutional approval. In the absence of any legal obligation, we asked the owners’ approval for feces collections and proceeding.

After the obtention of the verbal consent from animals owners, feces samples have been collected from nine different animal species including cat, dog, horse, sheep, rabbit, cow, pigs, goat, and donkey; from animals living in metropolitan France, more precisely in the Marseille metropolitan area, Southeastern France (Table 2). Dogs and cats were fed dry industrial dry kibble feed; horses were fed hay + straw + pellets; sheep and goats were fed pasture (grass) and dry supplementary feed; rabbits were fed dehydrated alfalfa + hay + pellets (other vegetables, cereals, mineral salts and vitamins); cows were fed hay + straw + pasture (grass) and whole plant maize silage; pigs were fed straw + dry pelleted feed (formula consisting mainly of maize, wheat, oats, peas, soybeans, cereals, oilseeds and minerals) and donkeys were fed grass and hay. Feces samples have been stored at +4°C for five weeks before being process for DNA extraction as reported below.

**Table 2.** Details of 407 feces samples here investigated for the presence of methanogens.

| Origin of samples | Collected number per sample | Collection sites |
| --- | --- | --- |
| Cat | 105 | Marseille |
| Dog | 52 | Marseille |
| Horse | 89 | Marseille and Carnoux |
| Sheep | 29 | Bourganeuf |
| Rabbit | 2 | Allauch |
| Cow | 57 | Bourganeuf and Allauch |
| Pig | 64 | Avignon |
| Goat | 5 | Allauch |
| Donkey | 4 | Allauch |

### DNA extraction and PCR assays

DNA extraction was performed by mixing 0.2 g of each feces sample with 500 μL of G2 buffer (QIAGEN, Hilden, Germany) in an Eppendorf tube (Fisher Scientific, Illkirch, France). Then, 0.3 g of acid-washed beads ≤ 106 μm (Sigma-Aldrich, Saint-Quentin Fallavier, France) were added in each tube and shake in a FastPrep BIO 101 device (MP Biomedicals, Illkirch, France) for 45 seconds for mechanical lysis before 10-minute incubation at 100°C. A 180 µL volume of the mixture was then incubated with 20 µL of proteinase K (QIAGEN) at 56°C overnight before a second mechanical lysis was performed. Total DNA was finally extracted with the EZ1 Advanced XL extraction kit (QIAGEN) and 200 μL eluted volume. Sterile phosphate-buffered saline (PBS) was used as a negative control in each DNA extraction run. Extracted DNA was incorporated into real-time PCR performed using Metha_16S_2_MBF: 5’-CGAACCGGATTAGATACCCG -3’ and Metha_16S_2_MBR: 5’-CCCGCCAATTCCTTTAAGTT-3’ primers (Eurogentec, Angers, France) and FAM_Metha_16S_2_MBP 6FAM-CCTGGGAAGTACGGTCGCAAG probe targeting the 16S DNA gene of methanogens, designed in our laboratory (Eurogentec). PCR amplification was done in a 20 μL volume including 15 μL of mix and 5 μL of extracted DNA. Five μL of ultra-pure water (Fisher Scientific) were used instead of DNA in the negative controls. The amplification reaction was performed in a CFX96 thermocycler (BioRad, Marnes-la-Coquette, France) incorporating a protocol with a cycle of 50°C for 2-minute, followed by 39 cycles of 95°C for 5-minute, 95°C for 5 seconds and finally 60°C for 30 seconds. Amplification of the archaeal 16S rRNA gene (primers used: SDArch0333aS15, 5-TCCAGGCCCTACGGG-3 and SDArch0958aA19, 5-YCCGGCGTTGAMTCCAATT-3) genes was performed as previously described [8, 9, 33, 34]. Sequencing reactions (Sangers’ method) were carried-out using the BigDye Terminator, version 1.1, cycle sequencing kit DNA according to the manufacturer’s instructions (Applied Biosystems, Foster City, USA). Nucleotide sequences were assembled using Chromas Pro software, version 1.7 (Technelysium Pty Ltd., Tewantin, Australia) and compared with sequences available in the GenBank database using the online NCBI BLAST program (http://blast.ncbi.nlm.nih.gov.gate1.inist.fr/Blast.cgi). We considered the sequences as belonging to the same species if the percentage of identity was >98.7%; as different species if between 95-98.7% and different genera if this threshold was < 95% with respect to the first hit obtained by BLAST (41).

### Multispacer sequence typing

We carried-out a multispacer sequence typing (MST) technique on all fecal specimens positive by PCR-sequencing as previously described in our laboratory (20,42). PCRs were realized in a 2720 Thermal Cycler (Applied Biosystems, Foster City, USA) and followed all the steps described for standard PCR used for the molecular analysis of fecal specimens. Negative controls consisting of PCR mixture without DNA template were included in each PCR run. All PCR products were sequenced in both directions using the same primers as used for PCRs in a 2720 Thermal Cycler (Applied Biosystems) with an initial 1-minute denaturation step at 96°C, followed by 25 cycles denaturation for 10 seconds each at 96°C, a 20 seconds annealing step at 50°C and a 4-minute extension step at 60°C. Sequencing products were purified using the MultiScreen 96-well plates Millipore (Merck, Molsheim, France), containing 5 % of Sephadex G-50 (Sigma-Aldrich), and sequences were analyzed on an ABI PRISM 31309 Genetic Analyzer (Applied Biosystem, Foster City, USA) and edited using the ChromasPro software (version 1.42; Technelysium Pty Ltd). For each intergenic spacer, a spacer type (ST) was defined as a sequence exhibiting unique genetic polymorphism (SNPs and indels). MST genotypes were defined as a unique combination of the four spacer sequences (20,42).

### Phylogenetic analyses

Sequences were edited using ChromasPro software (ChromasPro 1.7, Technelysium Pty Ltd., Tewantin, Australia). Molecular phylogenetic and evolutionary analyses were conducted in MEGA7 as previously described (43).

### Statistical analyses

We used R software for data analysis (https://www.r-project.org/). The Chi 2 test was used to compare the prevalence between the different animal species with a threshold α = 0.05.

## RESULTS

In this study, a total of 407 fecal specimens collected from nine different mammalian species were investigated by RT-PCR for the presence of methanogens (Table 2). Incorporating the 16S rRNA archaeal gene PCR primers newly designed in our laboratory into RT-PCR, we detected the presence of methanogen DNA in all mammals here investigated and none of the negative controls. 100.0% of cat feces specimens were positive with Ct values of 33.51 ± 1.28; 78.8% of dog feces specimens were positive with Ct values of 27.71 ± 0.94; 84.4% of horse feces specimens were positive with Ct values of 25 ± 2.95; 96.6% of sheep feces specimens were positive with Ct values of 27.19 ± 3.11; 100% of rabbit feces specimens were positive with Ct values of 27.1 ± 1.36; 100% of cow feces specimens were positive with Ct values of 24.11 ± 1.94; 100% of pig feces specimens were positive with Ct values of 22.15 ± 2.75; 80% of goat feces specimens were positive with Ct values of 19.18 ± 2.46 and 100% of donkey feces specimens were positive with Ct values of 18.82 ± 1.44 (Table 3 and Table 4). Prevalence as a function of RT-PCR differs significantly (p-value<0.05) between animal species (Table 4).

**Table 3.** Mean and standard deviation of the Ct values obtained in positive samples.

| Cat | Dog | Horse | Sheep | Rabbit | Cow | Pig | Goat | Donkey |
| --- | --- | --- | --- | --- | --- | --- | --- | --- |
| 33.51±1.28 | 27.71±0.94 | 25±2.95 | 27.19±3.11 | 27.1±1.36 | 24.11±1.94 | 22.15±2.75 | 19.18±2.46 | 18.82±1.44 |

**Table 4.** Comparison of prevalence based on RT-PCR between animal species.

| Animal species | Number of samples analyzed | Number of positive samples by RT-PCR | Prevalence [IC 95%] | p-value |
| --- | --- | --- | --- | --- |
| Cat | 105 | 105 | 100.0 [96.5-100.0] | 9.9X10<sup>-8</sup> |
| Dog | 52 | 41 | 78.8 [65.3-88.9] |  |
| Horse | 89 | 75 | 84.4 [75.0-91.1] |  |
| Sheep | 29 | 28 | 96.6 [82.2-99.9] |  |
| Rabbit | 2 | 2 | 100.0 [15.8-100.0] |  |
| Cow | 57 | 57 | 100.0 [93.7-100.0] |  |
| Pig | 64 | 64 | 100.0 [94.4-100.0] |  |
| Goat | 5 | 4 | 80.0 [28.4-99.5] |  |
| Donkey | 4 | 4 | 100.0 [39.8-100.0] |  |

Sequencing the PCR products was used to precise the identification of methanogens at the genus and species levels, in each sample. In cats, 50/105 successfully sequenced samples yielded 20 *Methanocorpusculum aggregans* (*M. aggregans*), 13 *Methanocorpusculum labreanum (M. labreanum)*, 09 *Methanobrevibacter millerae (M. millerae)*, 04 *M. smithii*, 02 *Methanobrevibacter thaueri (M. thaueri)*, and 02 *Methanobrevibacter olleyae (M. olleyae)*. In dogs, 30/52 successfully sequenced samples yielded 13 *M. labreanum*, 06 *M. smithii*, 05 *M. aggregans*, 03 *M. thaueri*, 02 *M. millerae*, and 01 *M. olleyae*. In horses, 24/89 successfully sequenced samples yielded 11 *M. aggregans*, 10 *M. olleyae*, 01 *M. smithii*, 01 *M. millerae*, and 01 *M. labreanum*. In sheep, 28/29 successfully sequenced samples yielded 23 *M. labreanum*, 03 *M. millerae*, 01 *M. smithii*, and 01 *M. aggregans*. In rabbits, 2/2 successfully sequenced samples yielded 02 *M. thaueri*. In cows, 44/57 successfully sequenced samples yielded 22 *M. aggregans*, 11 *M. millerae*, 06 *M. labreanum* and 05 *M. thaueri*. In pigs, 25/64 successfully sequenced samples yielded 12 *M. smithii*, 09 *M. millerae*, 03 *Methanomassiliicoccus luminiyensis* (*M. luminiyensis)* and 01 *M. olleyae*. In goats, 4/5 successfully sequenced samples yielded 03 *M. labreanum* and 01 *M. aggregans*. Finally, in donkeys, 4/4 successfully sequenced samples yielded 04 *M. aggregans* (Fig 1). So, we had 55/105 sequencing failures in cats; 11/41 in dogs; 51/75 in horses; 13/57 in cats and 39/64 in pigs. There were no sequencing failures in sheep, rats, goats, and donkeys (Table 5). Prevalence as a function of PCR-sequencing differs significantly (p-value<0.05) between animal species (Table 6).

**Table 5.** Number of sequencing failures observed in RT-PCR positive samples.

| Animal species | Number of positive samples by RT-PCR | Number of sequencing failures |
| --- | --- | --- |
| Cat | 105 | 55/105 |
| Dog | 41 | 11/41 |
| Horse | 75 | 51/75 |
| Sheep | 28 | 0/28 |
| Rabbit | 2 | 0/2 |
| Cow | 57 | 13/57 |
| Pig | 64 | 39/64 |
| Goat | 4 | 0/4 |
| Donkey | 4 | 0/4 |

**Table 6.**
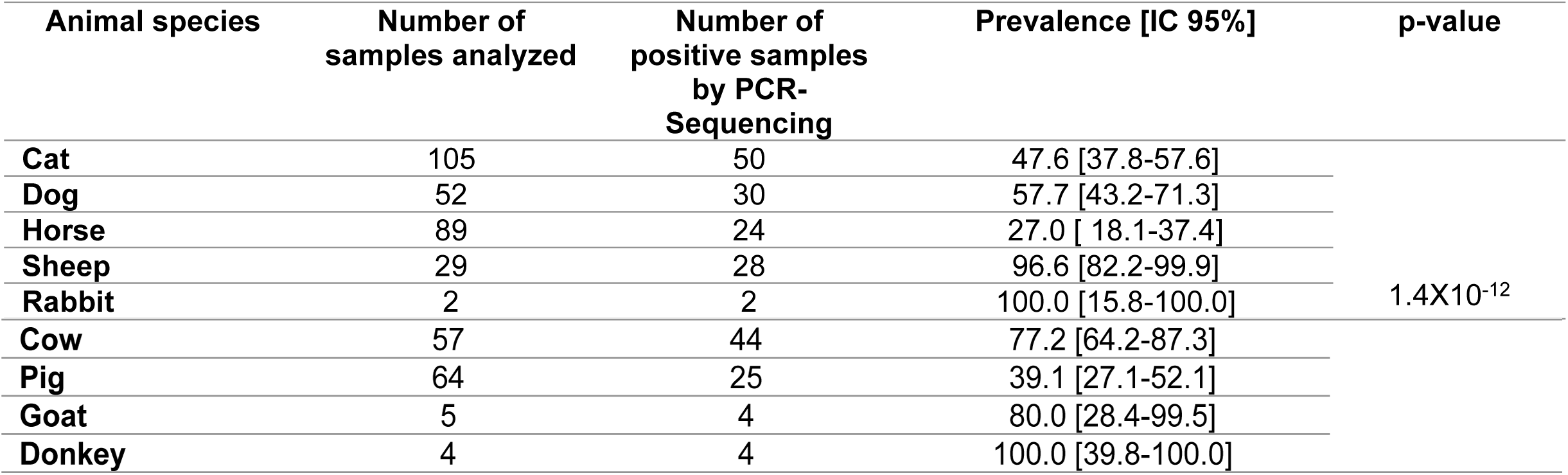
Comparison of prevalence based on PCR-sequencing between animal species.

| Animal species | Number of samples analyzed | Number of positive samples by PCR-Sequencing | Prevalence [IC 95%] | p-value |
| --- | --- | --- | --- | --- |
| Cat | 105 | 50 | 47.6 [37.8-57.6] | 1.4X10<sup>-12</sup> |
| Dog | 52 | 30 | 57.7 [43.2-71.3] |  |
| Horse | 89 | 24 | 27.0 [18.1-37.4] |  |
| Sheep | 29 | 28 | 96.6 [82.2-99.9] |  |
| Rabbit | 2 | 2 | 100.0 [15.8-100.0] |  |

|  |  |  |  |
| --- | --- | --- | --- |
| **Cow** | 57 | 44 | 77.2 [64.2-87.3] |
| **Pig** | 64 | 25 | 39.1 [27.1-52.1] |
| **Goat** | 5 | 4 | 80.0 [28.4-99.5] |
| **Donkey** | 4 | 4 | 100.0 [39.8-100.0] |

**Figure 1.**
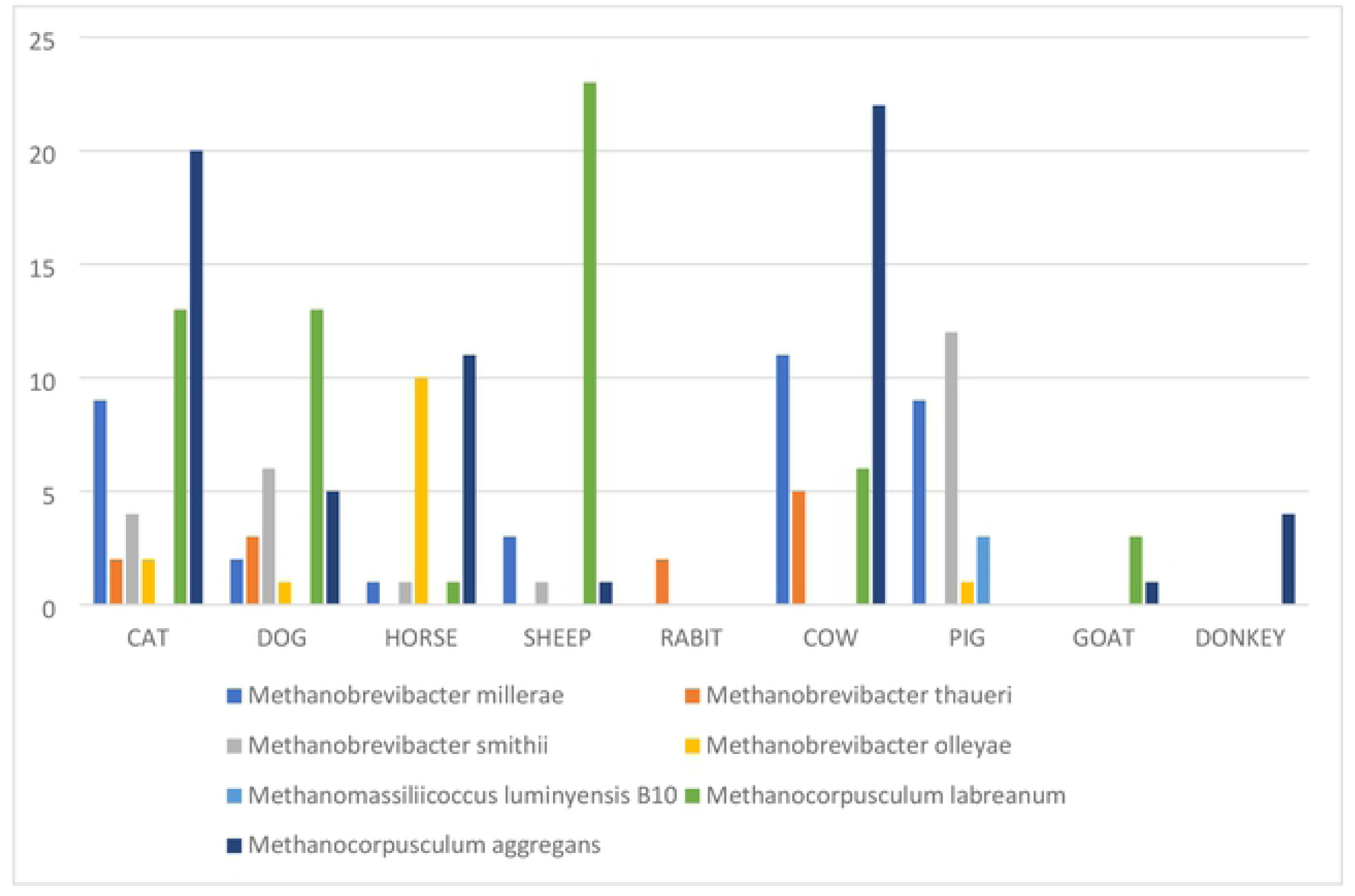
Species detected per sample according to their quantity.

We obtained a total of seven different species of methanogens in our study, of which three (*M. smithii, M. millerae* and *M. luminyensis*) are known to be part of the methanogens present in the human digestive tract (Fig 2 and Fig 3). The remaining four (*M. thaueri, M. olleyae, M. labreanum* and *M. aggregans*) are not known to date in humans. However, we did not find in our study the other ten species of methanogens present in the human digestive tract including *Methanobrevibacter arboriphilicus, M. oralis, Methanosphaera stadtmanae* (*M. stadtmanae), Candidatus* Methanomethylophilus alvus (*Ca*. Methanomethylophilus alvus), *Candidatus* Methanomassiliicoccus intestinalis (*Ca*. Methanomassiliicoccus intestinalis), *Methanoculleus chikugoensis* (*M. chikugoensis*), *Methanobacterium congolense* (*M. congolense*), *Methanoculleus bourgensis* (*M. bourgensis*), *Candidatus* Nitrososphaera evergladensis (*Ca*. Nitrososphaera evergladensis) and *Methanosarcinia mazei* (*M. mazei*) (Fig 2 and Fig 3).

**Figure 2.**
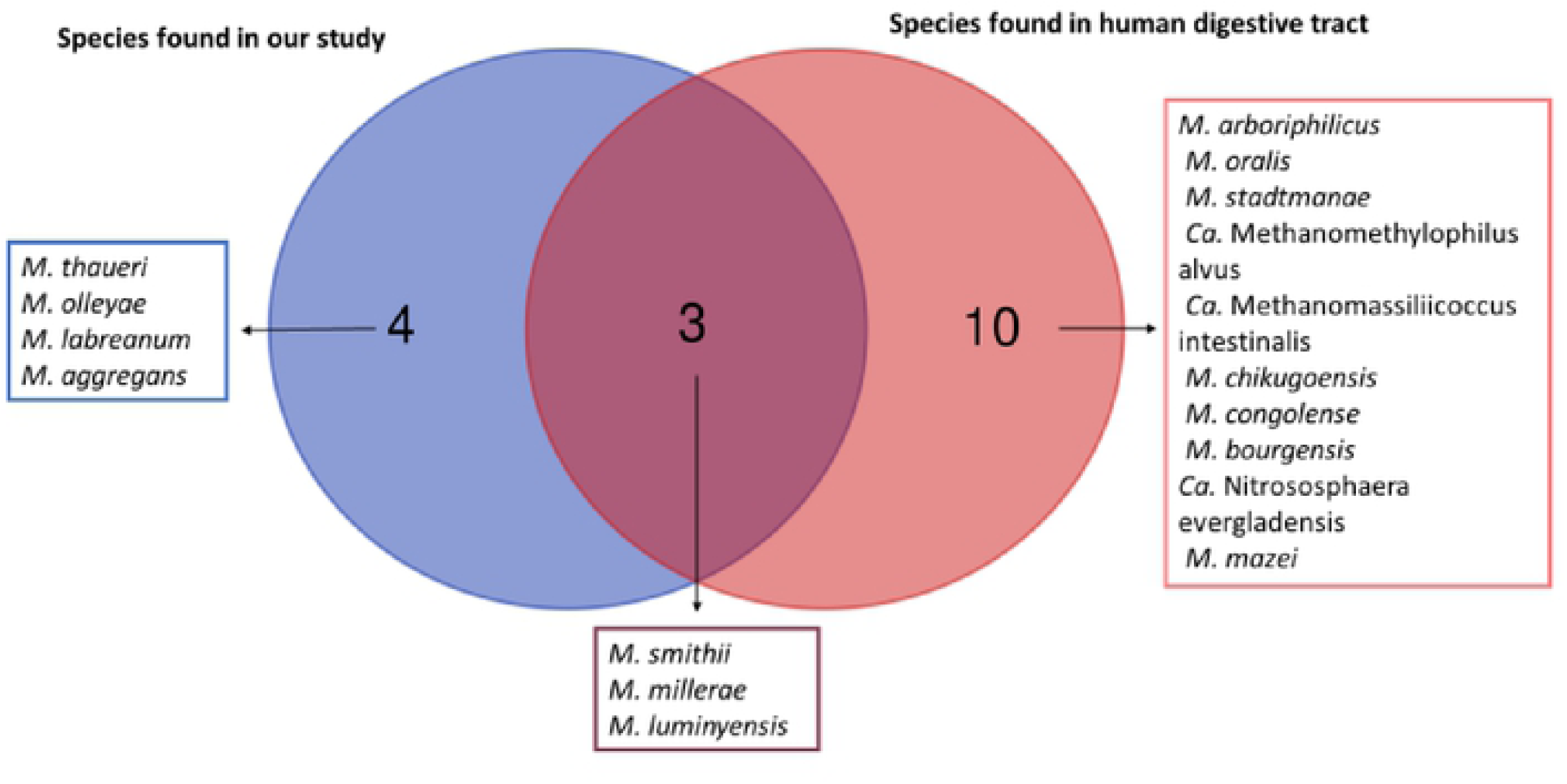
Venn diagram between the methanogens found in our study and those known from the human digestive tract.

**Figure 3.**
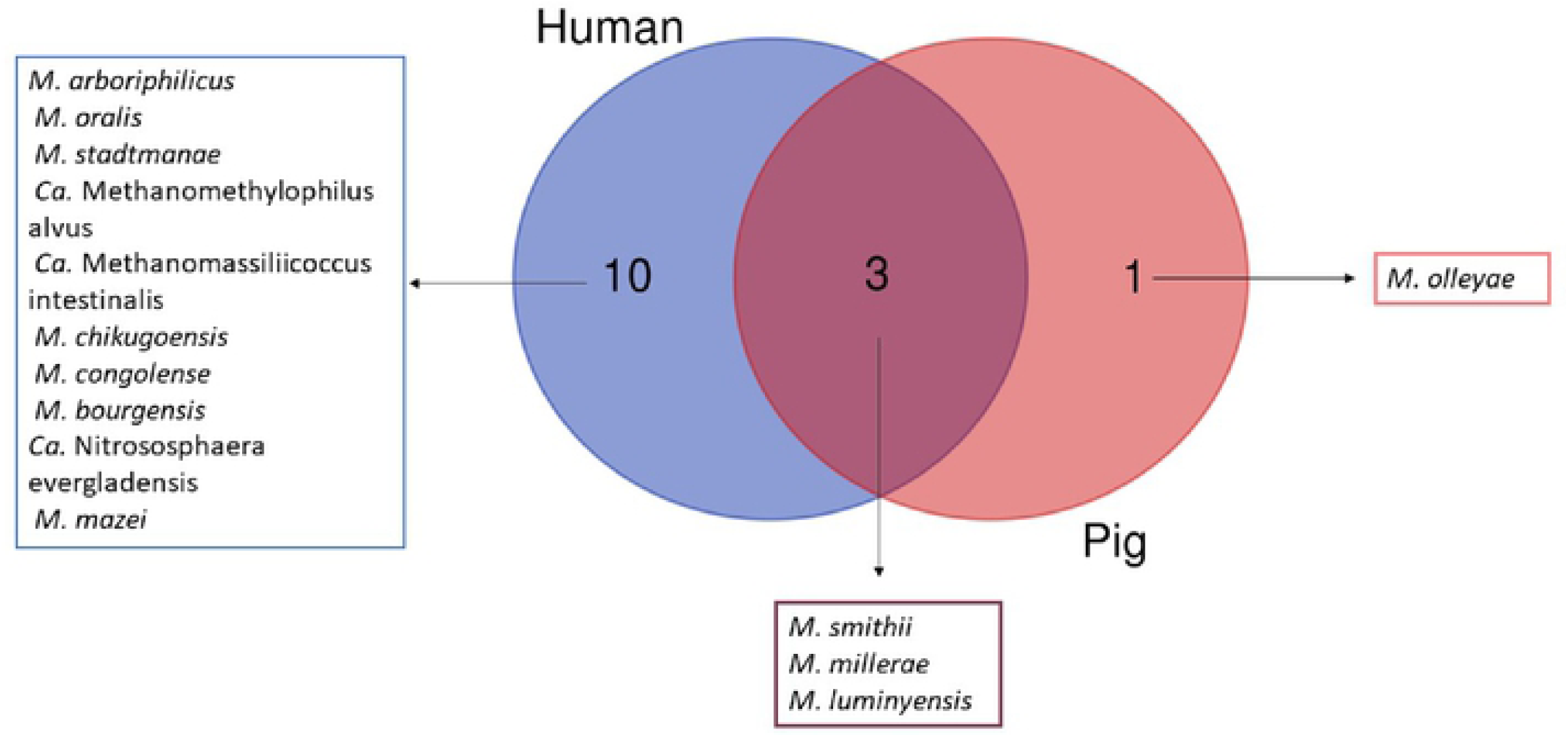
Venn diagram between the methanogens found in pigs and those known from the human digestive tract.

Among the 211 sequences obtained, 153 (72.51%) of them have an identity percentage greater than 99%; 43 (20.37%) have an identity percentage lower than 98.7% and 15 (7.10%) have an identity percentage lower than 95% (Table 7). The phylogenetic trees of sequences obtained with a percentage identity lower than 98.7% and sequences with a percentage identity lower than 95% indicated new species and new genera respectively (Fig 4 and Supplementary Figures). We obtained 24 *M. smithii* by PCR-sequencing including 12/24 (50%) in pigs, 6/24 (25%) in dogs, 4/24 (16.66%) in cats, and 1/24 (4.16%) in both sheep and horses (Table 8). Genotyping the 24 *M. smithii* revealed five different genotypes. Genotype 1 was found in 8/24 (33.33%); genotype 2 in 10/24 (41.66%); genotype 3 in 4/24 (16.66%); genotypes 4 and 5 in 1/24 (4.16%) each (Table 8).

**Table 7.** Percentage of identity among the sequences obtained.

| **Animal species** | **Percentage > 99%** | **Percentage < 98.7%** | **Percentage < 95%** |
| --- | --- | --- | --- |
| **Cat** | 50 | 0 | 0 |
| **Dog** | 30 | 0 | 0 |
| **Horse** | 7 | 15 | 2 |
| **Sheep** | 18 | 7 | 3 |
| **Rabbit** | 2 | 0 | 0 |
| **Cow** | 29 | 12 | 3 |
| **Pig** | 16 | 4 | 5 |
| **Goat** | 0 | 2 | 2 |
| **Donkey** | 1 | 3 | 0 |

**Table 8.** Summary of the results of multispacer sequence typing.

| **Samples** | **Origin** | **Collection sites** | **Genotype** |  |  |  |  |
| --- | --- | --- | --- | --- | --- | --- | --- |
|  |  |  | **Spacer1** | **Spacer2** | **Spacer 3** | **Spacer 4** | **Spacer type\*** |
| **1** | Sheep | Bourganeuf |  |  | X |  | 1 |
| **2** | Horse | Marseille |  |  | X |  | 1 |
| **3** | Pig | Avignon | X | X | X | X | 2 |
| **4** | Pig | Avignon | X | X | X | X | 2 |
| **5** | Pig | Avignon | X | X | X | X | 2 |
| **6** | Pig | Avignon | X | X | X | X | 2 |
| **7** | Pig | Avignon | X | X | X |  | 3 |
| **8** | Pig | Avignon | X | X |  |  | 4 |
| **9** | Pig | Avignon |  | X | X |  | 5 |
| **10** | Pig | Avignon |  |  | X |  | 1 |
| **11** | Pig | Avignon | X | X | X | X | 2 |
| **12** | Pig | Avignon | X | X | X |  | 3 |
| **13** | Pig | Avignon | X | X | X | X | 2 |
| **14** | Pig | Avignon | X | X | X | X | 2 |
| **15** | Dog | Marseille | X | X | X | X | 2 |
| **16** | Dog | Marseille | X | X | X | X | 2 |
| **17** | Dog | Marseille | X | X | X |  | 3 |
| **18** | Dog | Marseille | X | X | X |  | 3 |
| **19** | Dog | Marseille | X | X | X | X | 2 |
| **20** | Dog | Marseille |  |  | X |  | 1 |
| **21** | Cat | Marseille |  |  | X |  | 1 |
| **22** | Cat | Marseille |  |  | X |  | 1 |
| **23** | Cat | Marseille |  |  | X |  | 1 |
| **24** | Cat | Marseille |  |  | X |  | 1 |

**Figure 4.**
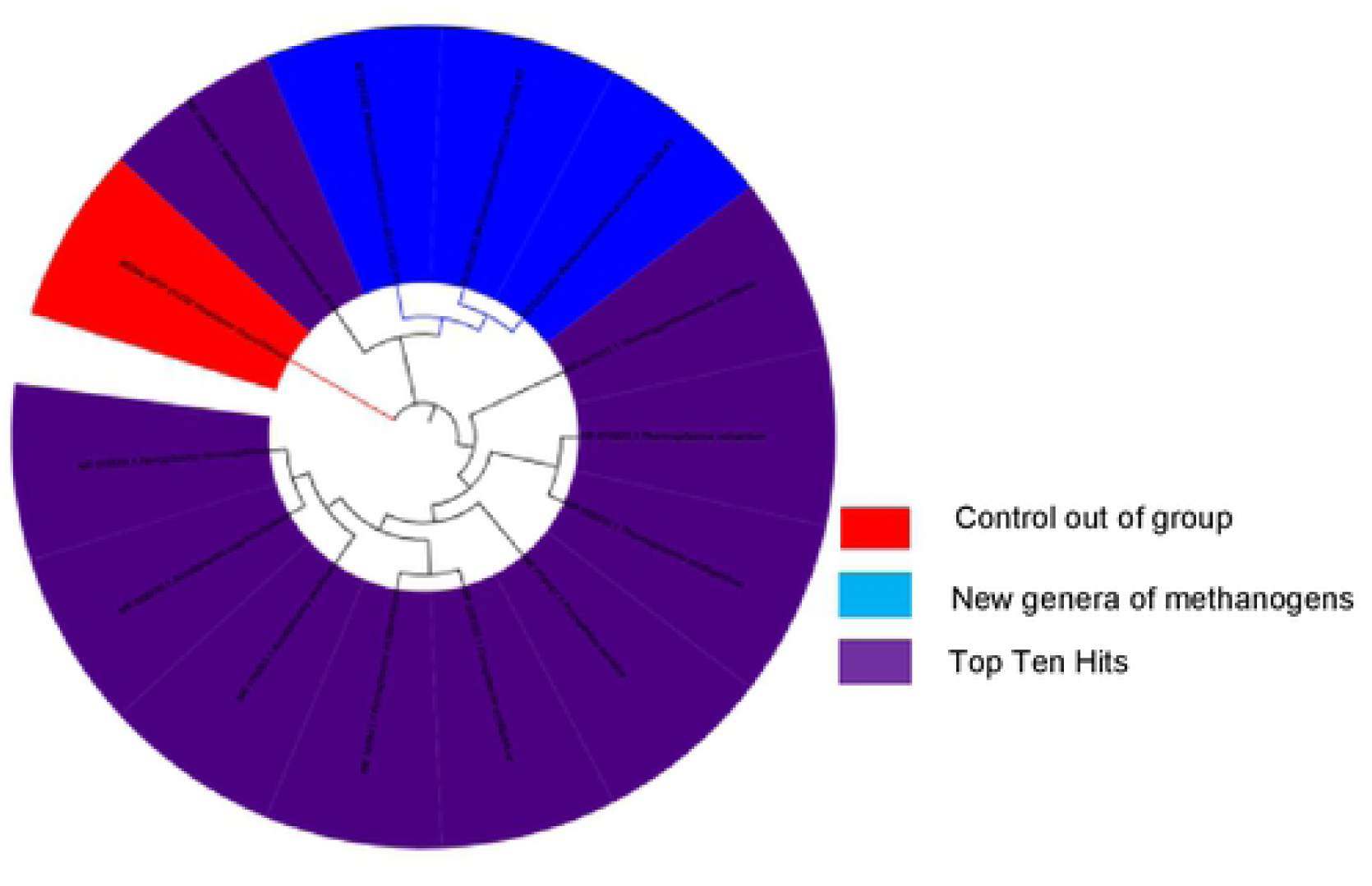
Molecular phylogenetic analysis, based on 16S rRNA partial gene, showed the position of *Methanomassiliicoccus-like* sqeuences detected in feces of pig. The evolutionary history was inferred using the Neighbor-Joining method. The optimal tree with the sum of branch length= 0.82722721 is shown. The percentage of replicate trees in which the associated taxa clustered together in the bootstrap test (1.000 replicates) are shown next to the branches. The tree is drawn to scale, with branch lengths in the same units as those of the evolutionary distances used to infer the phylogenetic tree. The evolutionary distances were computed using the Maximum Composite Likelihood method and are in the units of the number of base substitutions per site. The analysis involved 14 nucleotide sequences. All positions containing gaps and missing data were eliminated. There was a total of 415 positions in the final dataset. Evolutionary analyses were conducted in MEGA7.

## DISCUSSION

It is known and published that methanogens colonize the gastrointestinal tract of certain mammals, particularly herbivorous ones [35]. Most methanogens identified in mammals belong to the phylum Euryarchaeota, with a high percentage of the species *M. smithii* [36], a species being the most prevalent one in humans (44). Our report is the largest one showing the presence of methanogens in nine mammals in the same study. Our results confirmed the published data on the presence of methanogens in the digestive tract of cats, dogs, horses, cows, sheep, rabbits, goats, pigs, and donkeys (18,19,25–31,33–35). In addition, all methanogens found in this study belong to the phylum Euryarchaeota, which is in accordance with the results obtained in studies conducted on the human digestive tract (45,46). Our results give an insight on the concentration of methanogens present in the intestinal microbiota of each animal species analyzed and on the prevalence of methanogens in domestic animals by humans.

The results of the analysis of the 16S RNA sequences obtained from our samples show that there is a real diversity of methanogenic archaea genera (*Methanosphaera, Methanocorpusculum, Methanocalculus, Methanoculleus, Methanogenium, Methanoplanus, Methanolacinia, Methanobacterium, Methanomicrobium, Methanomassiliicoccus* and *Methanobrevibacter*) in the digestive tract of animals (cats, dogs, horses, sheep, cows, rabbits, goats, pigs and donkeys) as in humans (45,46). All sequences with a percentage lower than 98.7% have been deposited in the GenBank database (accession no MT587812 to MT587864) and EBI database (accession no MT793590; MT819603; MT822292; MT822293; and MT822482).

This study demonstrated for the first time the presence of the genus *Methanomassiliicoccus* in animals, specifically *M. luminiyensis* in pigs. This methanogen was until now known only in humans (47). This could be explained by the fact that the pig is an omnivore which means that its diet is very close to that of humans compared to other animals. However, the only time this species was isolated and cultured in humans was in a healthy 86-year-old Caucasian man who most probably consumed pig (47). This suggested a probable link between consumption of pig meat and/or contact with pig and the presence of *M. luminiyensis* in humans. In addition, 50% of *M. smithii* in our study were found in pigs, indicating that *M. smithii* was the most prevalent methanogen in the digestive tract of pigs, consistent with work carried out in humans where the high prevalence of *M. smithii* in the digestive tract has been demonstrated (44,46).

These results are fairly representative of the methanogen community present in the digestive tract of certain animals domesticated by humans and other future studies must be done to try to cultivate methanogens here detected by molecular biology to better understand the dynamics of methanogens in animals and also the likely acquisition of methanogens in humans through direct contact with these animals or through consumption of the meat and/or milk of certain animals, in particular cows, since a recent study has demonstrated an association between the acquisition of *M. smithii* in children and the consumption of dairy products (48).

## ACKNOWLEDGEMENTS

This work was supported by a grant of GDR Archaea, Centre National de la Recherche Scientifique, Paris, France.

## SUPPLEMENTARY FIGURES

**Supplementary Figure 1.**
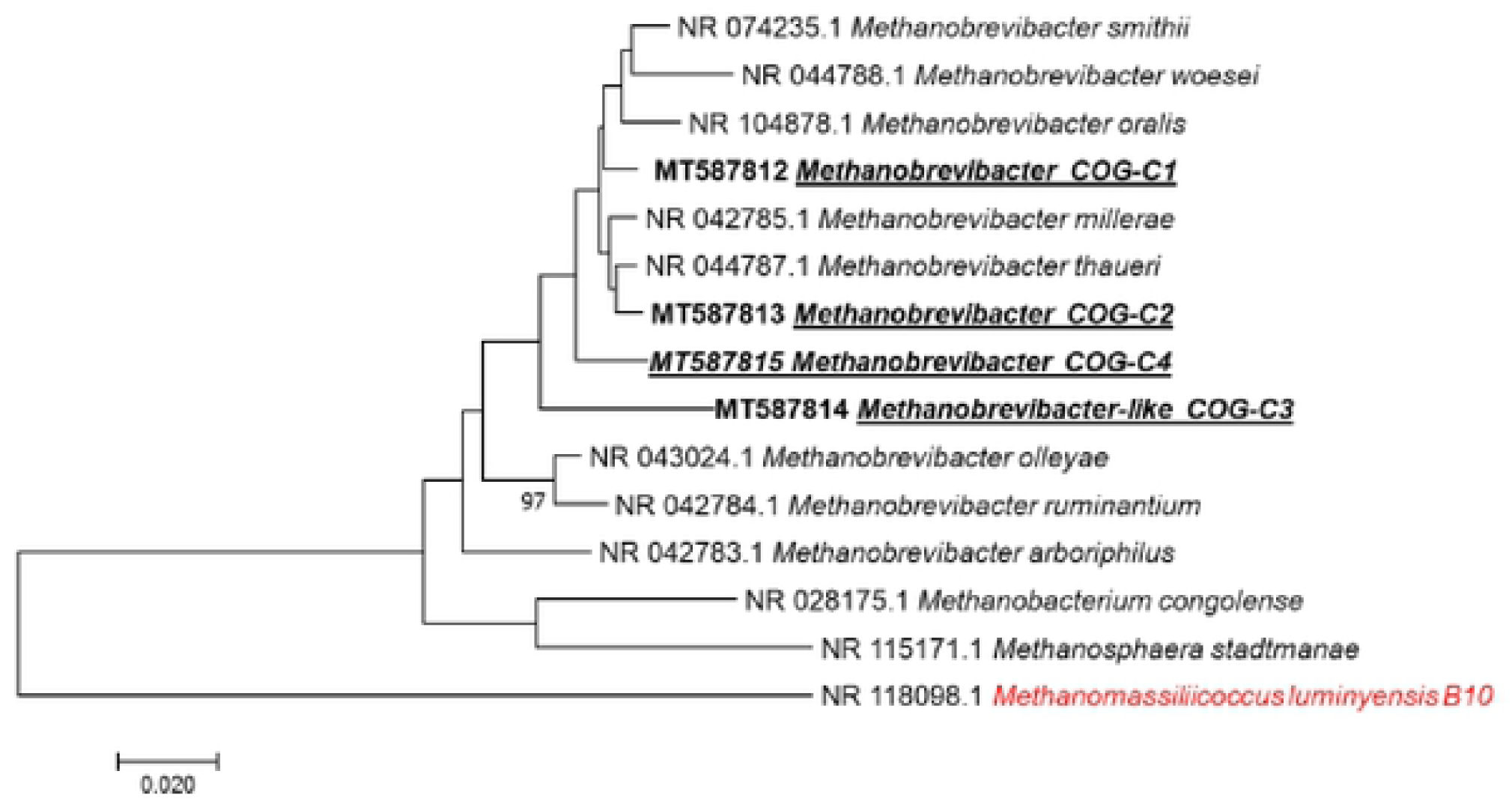
Molecular phylogenetic analysis, based on 16S rRNA partial gene, showed the position of *Methanobrevibacter* sequences detected in feces of cow. The evolutionary history was inferred using the Neighbor-Joining method. The optimal tree with the sum of branch length =0.56972409 is shown. The percentage of replicate trees in which the associated taxa clustered together in the bootstrap test (1.000 replicates) are shown next to the branches. The tree is drawn to scale, with branch lengths in the same units as those of the evolutionary distances used to infer the phylogenetic tree. The evolutionary distances were computed using the Maximum Composite Likelihood method and are in the units of the number of base substitutions per site. The analysis involved 15 nucleotide sequences. All positions containing gaps and missing data were eliminated. There was a total of 447 positions in the final dataset. Evolutionary analyses were conducted in MEGA7. Bootstrap values ≥95%, are indicated at nodes.

**Supplementary Figure 2.**
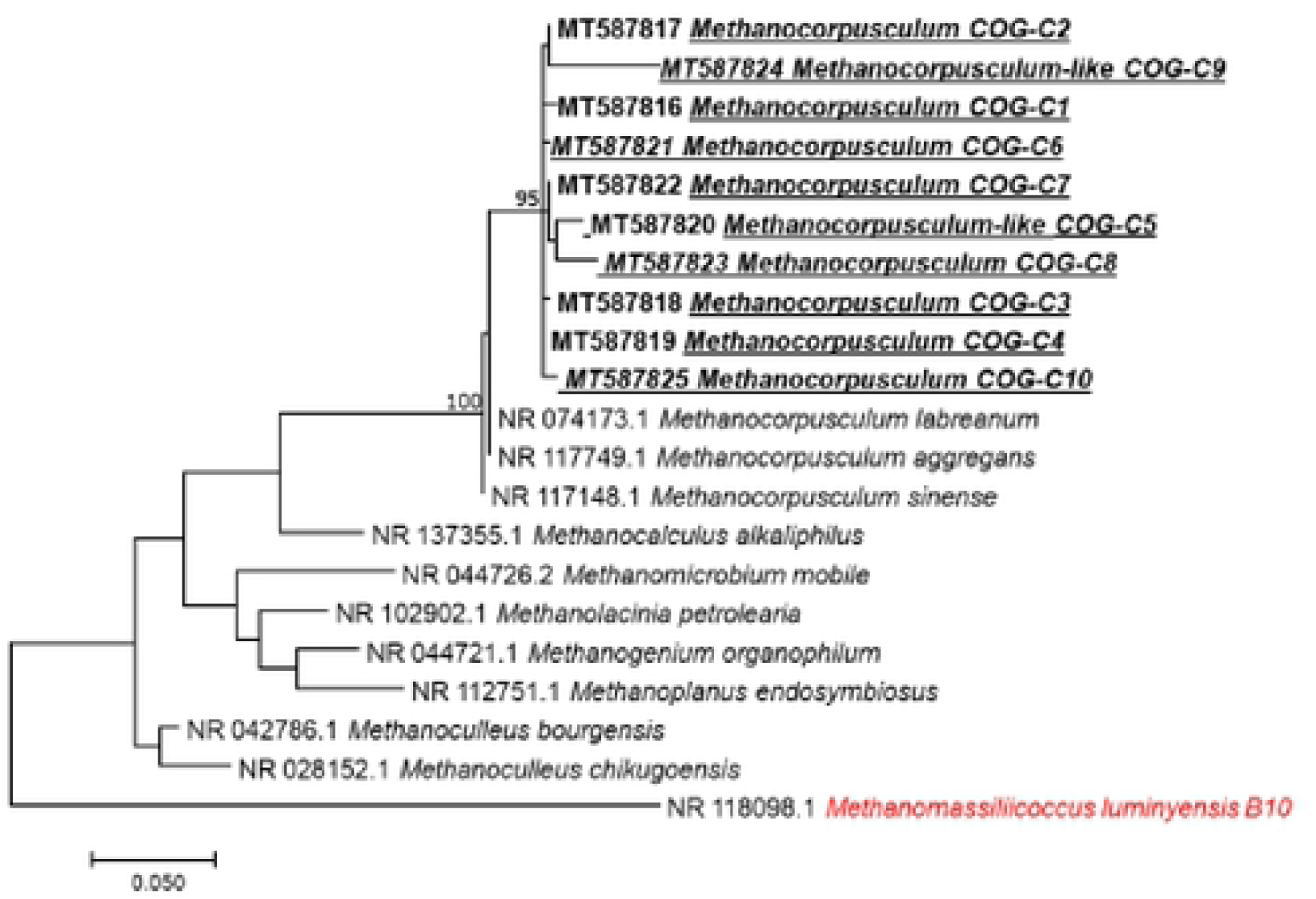
Molecular phylogenetic analysis, based on 16S rRNA partial gene, showed the position of *Methanocorpusculum* sequences detected in feces of cow. The evolutionary history was inferred by using the Maximum Likelihood method based on the Kimura 2-parameter model. The tree with the highest log likelihood (−1869.71) is shown. The percentage of trees in which the associated taxa clustered together is shown next to the branches. Initial tree(s) for the heuristic search were obtained automatically by applying Neighbor-Join and BioNJ algorithms to a matrix of pairwise distances estimated using the Maximum Composite Likelihood (MCL) approach, and then selecting the topology with superior log likelihood value. The tree is drawn to scale, with branch lengths measured in the number of substitutions per site. The analysis involved 21 nucleotide sequences. All positions containing gaps and missing data were eliminated. There was a total of 375 positions in the final dataset. Evolutionary analyses were conducted in MEGA7. Bootstrap values ≥95%, are indicated at nodes.

**Supplementary Figure 3.**
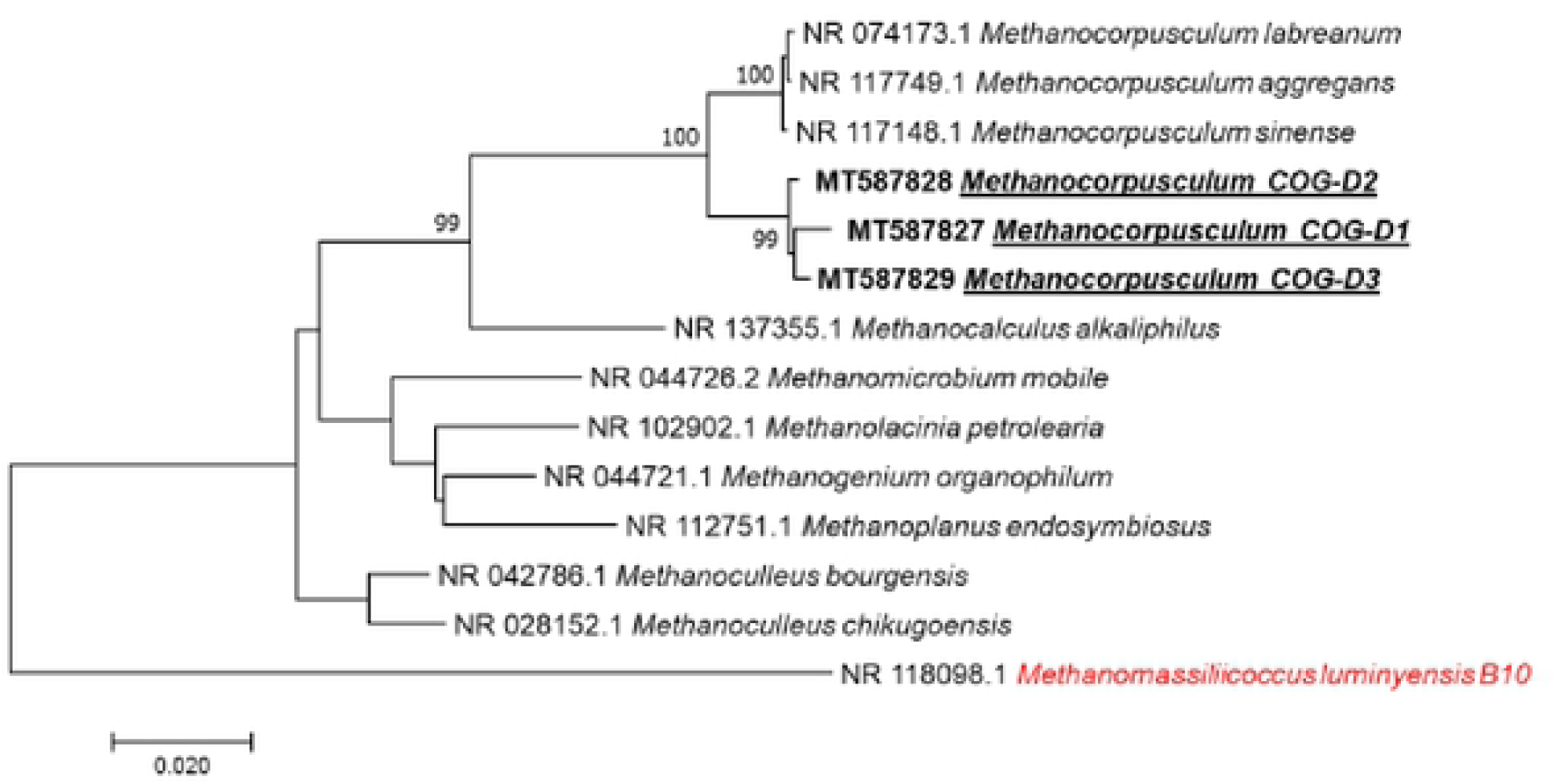
Molecular phylogenetic analysis,based on 16S rRNA partial gene, showed the position of *Methanocorpusculum* sequences detected in feces of donkey. The evolutionary history was inferred using the Neighbor-Joining method. The optimal tree with the sum of branch length=0.41522849 is shown. The percentage of replicate trees in which the associated taxa clustered together in the bootstrap test(1.000 replicates)are shown next to the branches. The tree is drawn to scale,with branch lengths in the same units as those of the evolutionary distances used to infer the phylogenetic tree. The evolutionary distances were computed using the Maximum Composite Likelihood method and are in the units of the number of base substitutions per site. The analysis involved 14 nucleotide sequences. All positions containing gaps and missing data were eliminated. There was a total of 476 positions in the final dataset. Evolutionary analyses were conducted in MEGA7. Bootstrap values ≥95% are indicated at nodes.

**Supplementary Figure 4.**
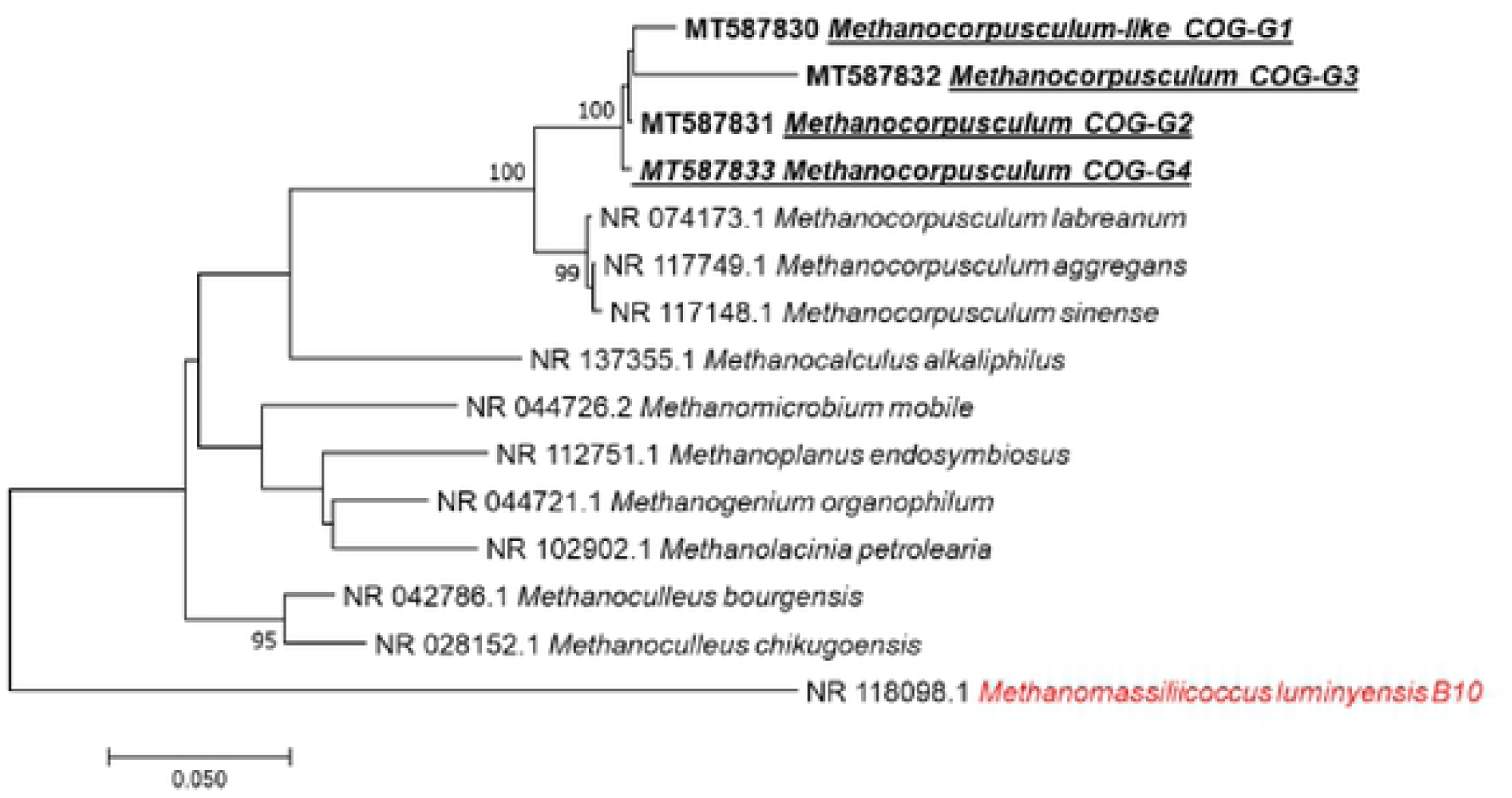
Molecular phylogenetic analysis, based on 16S rRNA partial gene, showed the position of *Methanocorpusculum* sequences detected in feces of goat. The evolutionary history was inferred using the Neighbor-Joining method. The optimal tree with the sum of branch length= 0.79665377 is shown. The percentage of replicate trees in which the associated taxa clustered together in the bootstrap test (1.000 replicates) are shown next to the branches. The tree is drawn to scale, with branch lengths in the same units as those of the evolutionary distances used to infer the phylogenetic tree. The evolutionary distances were computed using the Maximum Composite Likelihood method and are in the units of the number of base substitutions per site. The analysis involved 15 nucleotide sequences. All positions containing gaps and missing data were eliminated. There was a total of 427 positions in the final dataset. Evolutionary analyses were conducted in MEGA7. Bootstrap values ≥95%, are indicated at nodes.

**Supplementary Figure 5.**
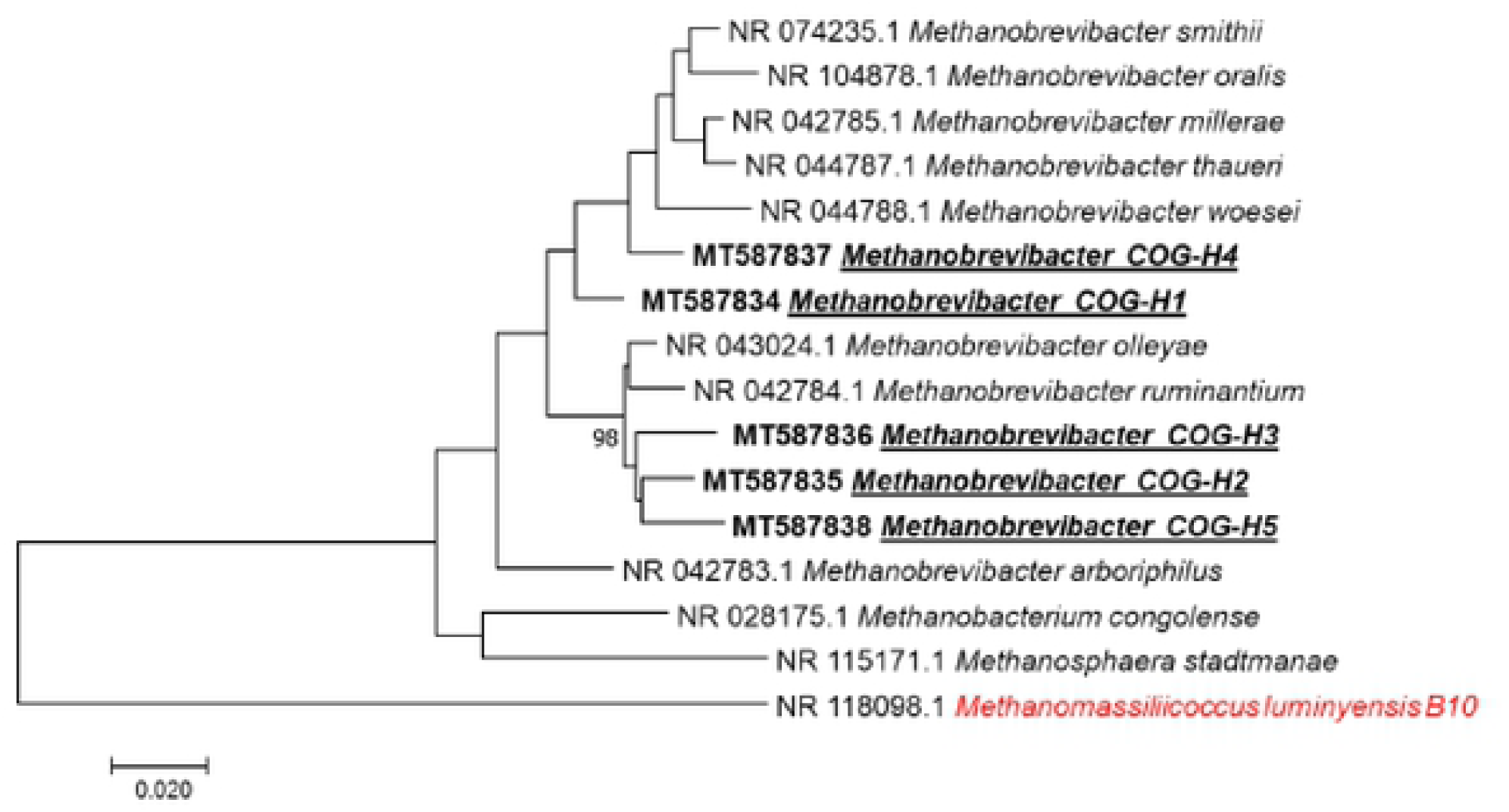
Molecular phylogenetic analysis, based on 16S rRNA partial gene, showed the position of *Methanobrevibacter* sequences detected in feces of horse. The evolutionary history was inferred using the Neighbor-Joining method. The optimal tree with the sum of branch length = 0.58360989 is shown. The percentage of replicate trees in which the associated taxa clustered together in the bootstrap test (1.000 replicates) are shown next to the branches. The tree is drawn to scale, with branch lengths in the same units as those of the evolutionary distances used to infer the phylogenetic tree. The evolutionary distances were computed using the Maximum Composite Likelihood method and are in the units of the number of base substitutions per site. The analysis involved 16 nucleotide sequences. All positions containing gaps and missing data were eliminated. There was a total of 408 positions in the final dataset. Evolutionary analyses were conducted in MEGA7. Bootstrap values ≥95%, are indicated at nodes.

**Supplementary Figure 6.**
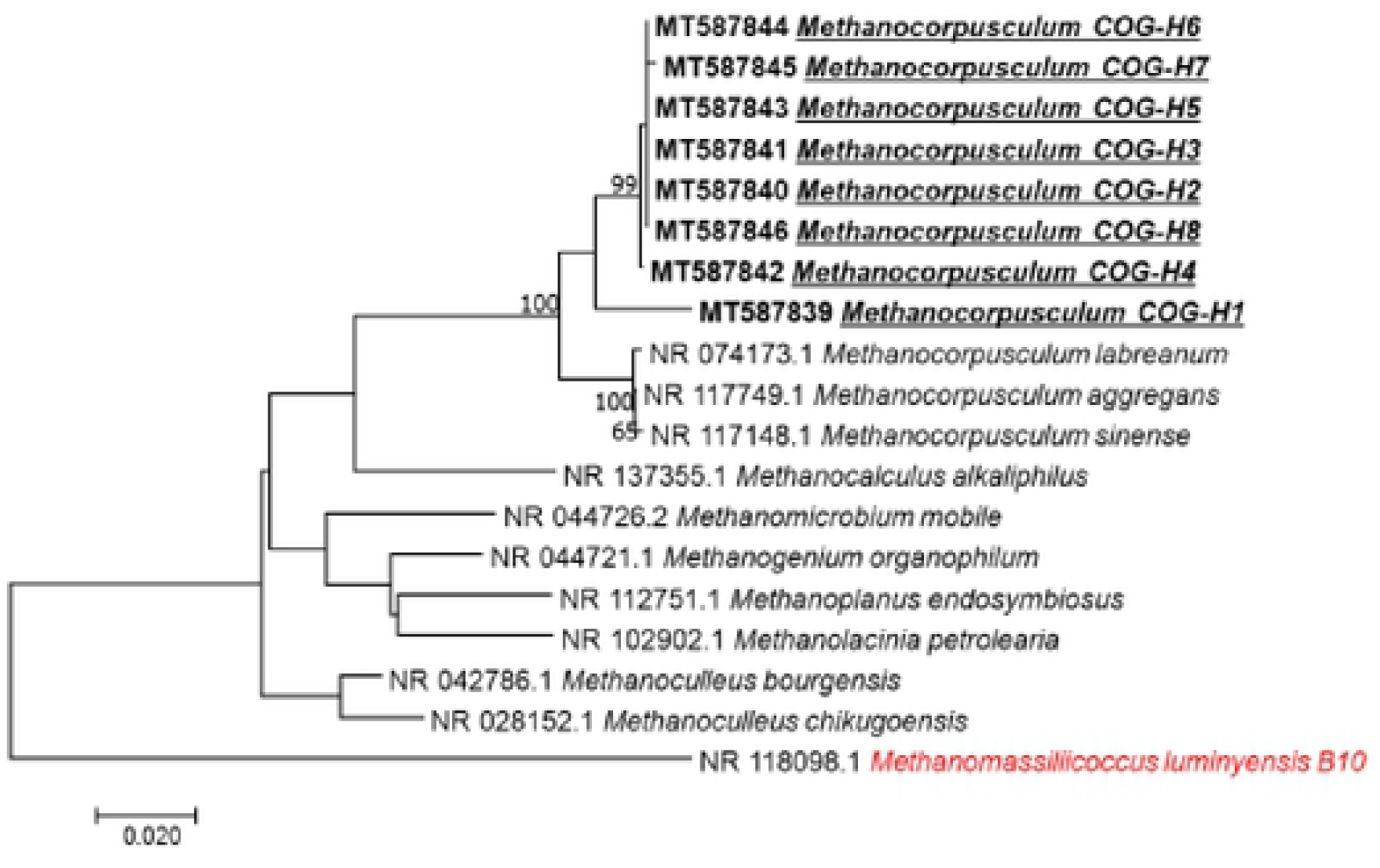
Molecular phylogenetic analysis, based on 16S rRNA partial gene, showed the position of *Methanocorpusculum* sequences detected in feces of horse. The evolutionary history was inferred using the Neighbor-Joining method. The optimal tree with the sum of branch length= 0.52514892 is shown. The percentage of replicate trees in which the associated taxa clustered together in the bootstrap test (1.000 replicates) are shown next to the branches. The tree is drawn to scale, with branch lengths in the same units as those of the evolutionary distances used to infer the phylogenetic tree. The evolutionary distances were computed using the Maximum Composite Likelihood method and are in the units of the number of base substitutions per site. The analysis involved 19 nucleotide sequences. All positions containing gaps and missing data were eliminated. There was a total of 417 positions in the final dataset. Evolutionary analyses were conducted in MEGA7. Bootstrap values ≥95%, are indicated at nodes.

**Supplementary Figure 7.**
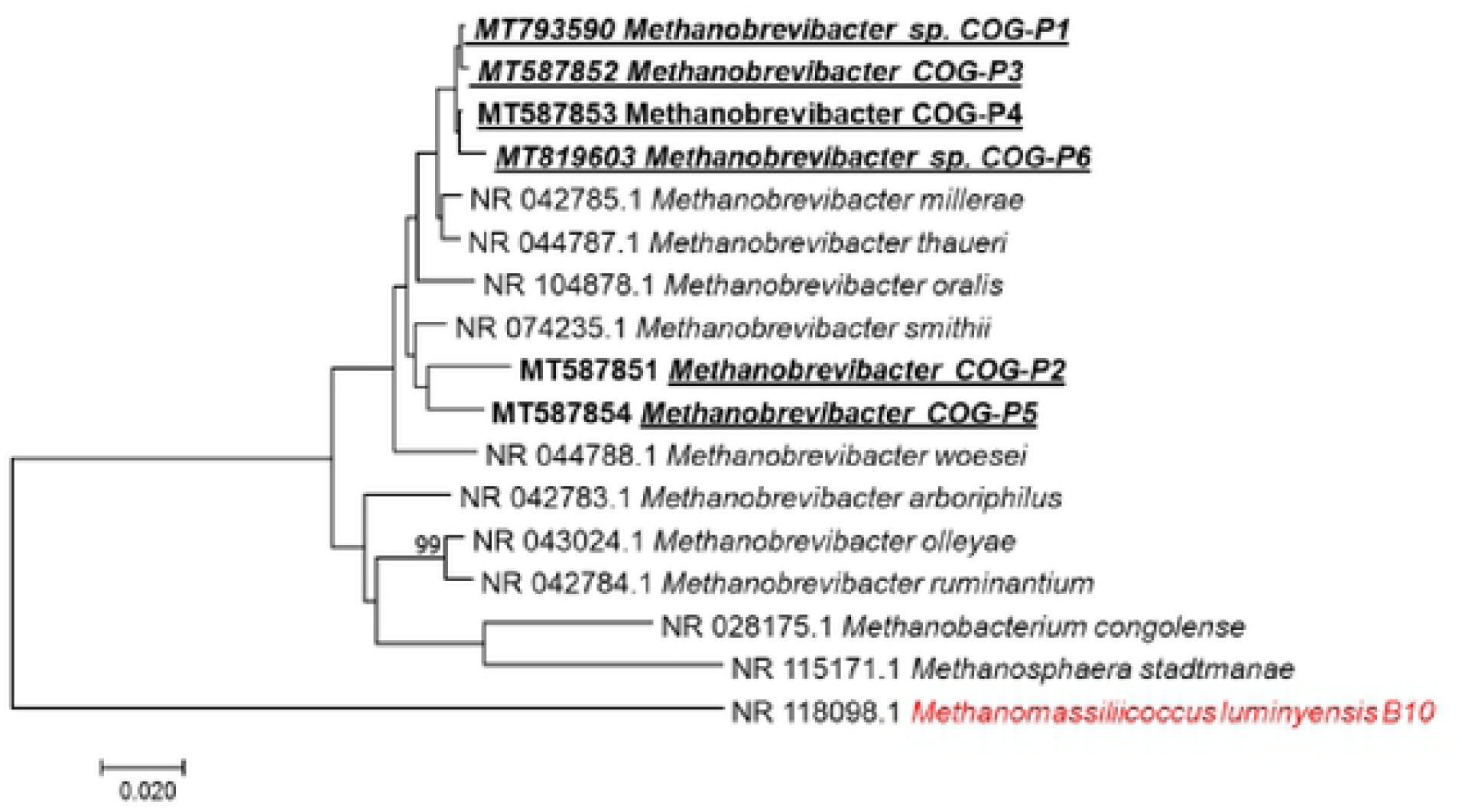
Molecular phylogenetic analysis, based on 16S rRNA partial gene, showed the position of *Methanobrevibacter* sequences detected in feces of pig. The evolutionary history was inferred using the Neighbor-Joining method. The optimal tree with the sum of branch length= 0.56411513 is shown. The percentage of replicate trees in which the associated taxa clustered together in the bootstrap test (1.000 replicates) are shown next to the branches. The tree is drawn to scale, with branch lengths in the same units as those of the evolutionary distances used to infer the phylogenetic tree. The evolutionary distances were computed using the Maximum Composite Likelihood method and are in the units of the number of base substitutions per site. The analysis involved 17 nucleotide sequences. All positions containing gaps and missing data were eliminated. There was a total of 467 positions in the final dataset. Evolutionary analyses were conducted in MEGA7. Bootstrap values ≥95%, are indicated at nodes.

**Supplementary Figure 8.**
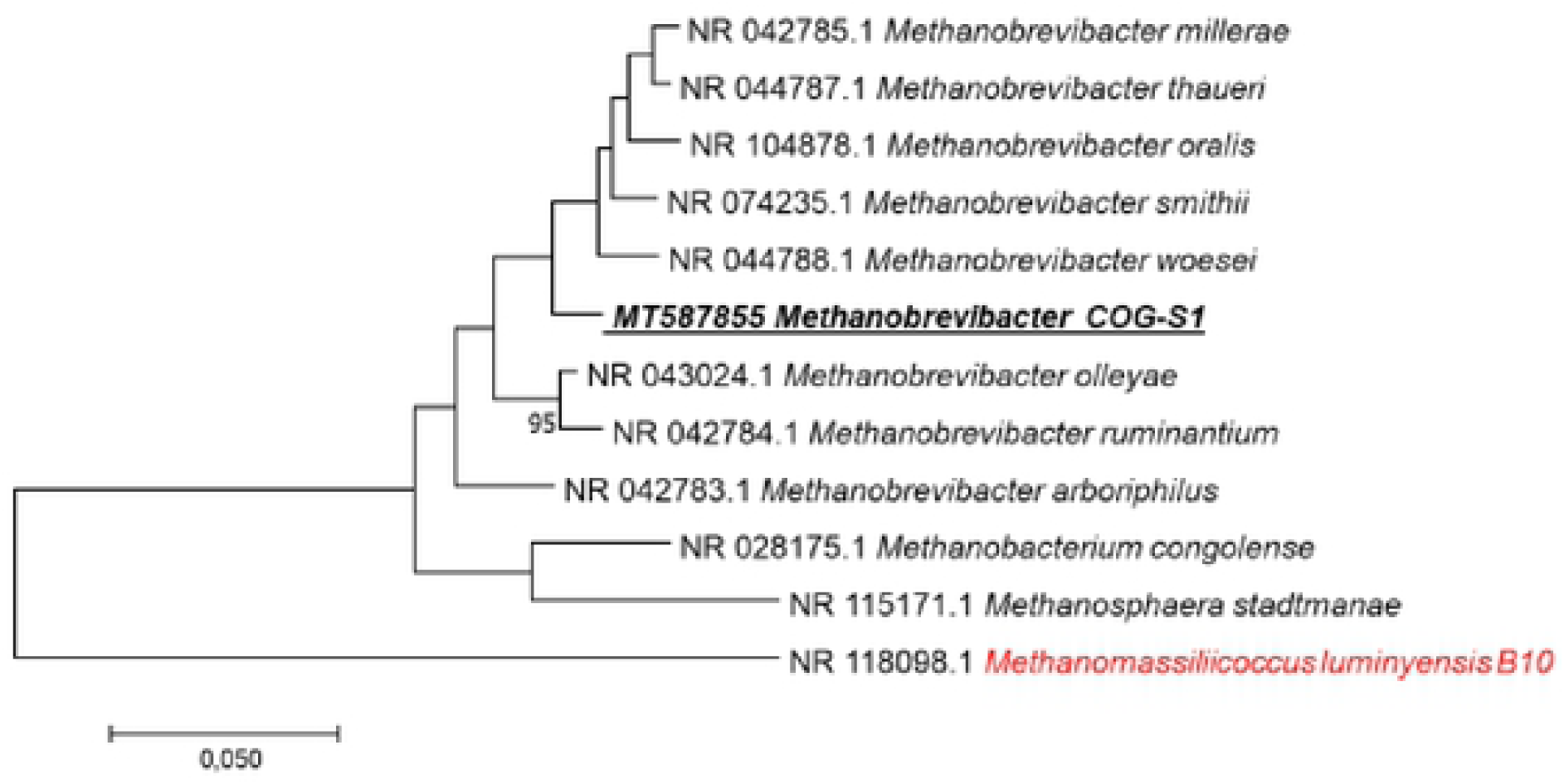
Molecular phylogenetic analysis, based on 16S rRNA partial gene, showed the position of *Methanobrevibacter* sequence detected in feces of sheep. The evolutionary history was inferred by using the Maximum Likelihood method based on the Kimura 2-parameter model. The tree with the highest log likelihood (−1972.37) is shown. The percentage of trees in which the associated taxa clustered together is shown next to the branches. Initial tree(s) for the heuristic search were obtained automatically by applying Neighbor-Join and BioNJ algorithms to a matrix of pairwise distances estimated using the Maximum Composite Likelihood (MCL) approach, and then selecting the topology with superior log likelihood value. The tree is drawn to scale, with branch lengths measured in the number of substitutions per site. The analysis involved 12 nucleotide sequences. All positions containing gaps and missing data were eliminated. There was a total of 554 positions in the final dataset. Evolutionary analyses were conducted in MEGA7. Bootstrap values 95%, are indicated at nodes.

**Supplementary Figure 9.**
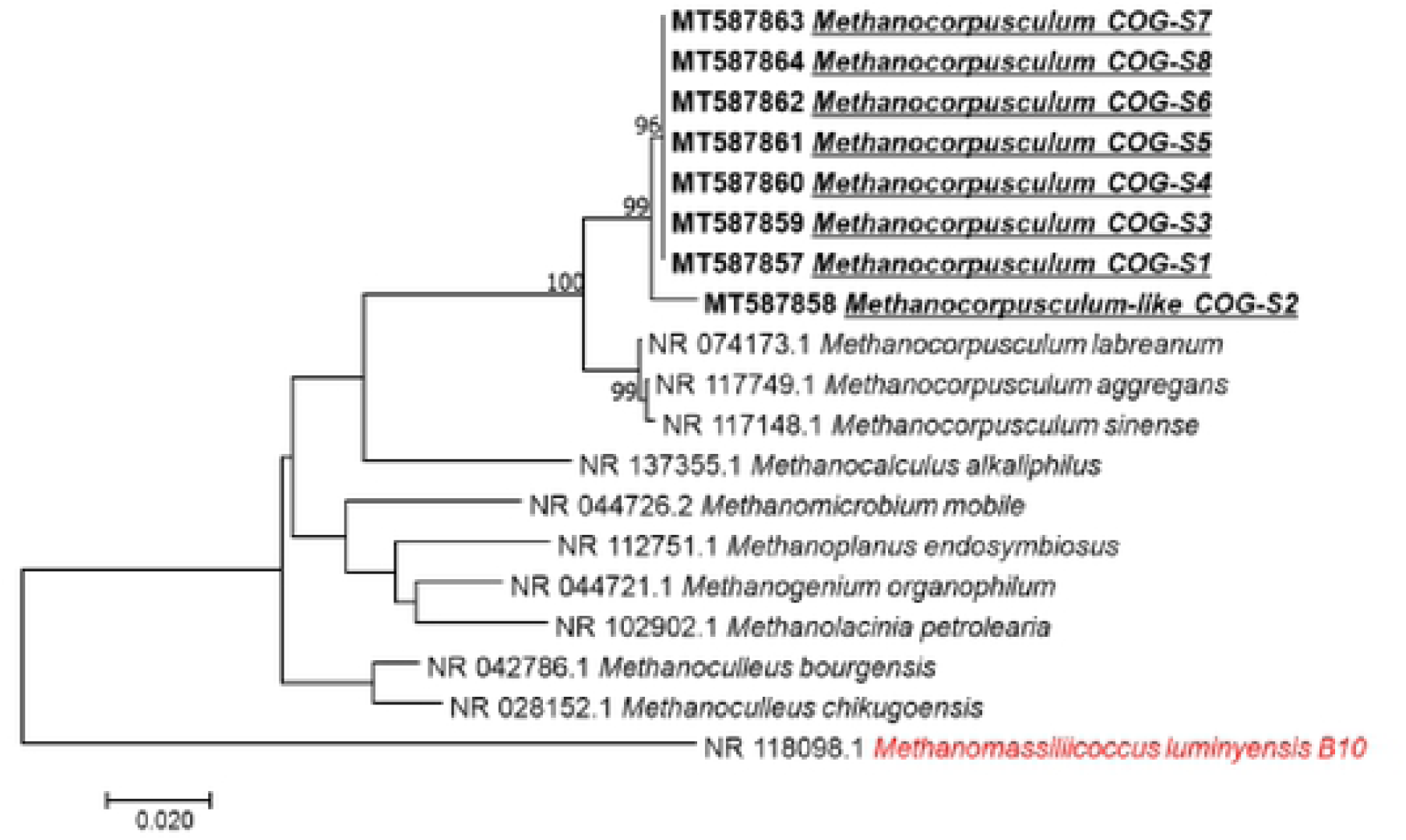
Molecular phylogenetic analysis, based on 16S rRNA partial gene, showed the position of *Methanocorpusculum* sequences detected in feces of sheep. The evolutionary history was inferred using the Neighbor-Joining method. The optimal tree with the sum of branch length= 0.48477167 is shown. The percentage of replicate trees in which the associated taxa clustered together in the bootstrap test (1.000 replicates) are shown next to the branches. The tree is drawn to scale, with branch lengths in the same units as those of the evolutionary distances used to infer the phylogenetic tree. The evolutionary distances were computed using the Maximum Composite Likelihood method and are in the units of the number of base substitutions per site. The analysis involved 19 nucleotide sequences. All positions containing gaps and missing data were eliminated. There was a total of 441 positions in the final dataset. Evolutionary analyses were conducted in MEGA7. Bootstrap values ≥95% are indicated at nodes.

